# Co-culture of *Saccharomyces cerevisiae* (VS3) and *Pichia stipitis* (NCIM 3498) for bioethanol production using concentrated *Prosopis juliflora* acid hydrolysate

**DOI:** 10.1101/601278

**Authors:** Shaik Naseeruddin, Suseelendra Desai, L Venkateswar Rao

## Abstract

Bioethanol production from lignocellulosic biomass is a viable option for improving energy security and reducing green house emissions. In the present study *Prosopis juliflora*, an invasive tree found through out India, with total carbohydrate content of 67.4 +/- 6% was selected as lignocellulosic feedstock for bioethanol production. The hydrolysate obtained after biphasic dilute acid hydrolysis contained initial sugar concentration of 18.70 +/- 0.16 g/L and hence to increase the ethanol yield it was concentrated to 33.59 +/- 0.52 g/L (about two-folds) by vacuum distillation. The concentration of sugars, phenols and furans was analyzed before and after concentration process. The concentrated hydrolysate was further detoxified by over liming, neutralization and charcoal treatment and later used for ethanol fermentation by mono- and co culture method. Monoculture of *Saccharomyces cerevisiae* (VS3) and *Pichia stipitis* (NCIM 3498) produced 8.52 +/- 0.43 and 4.52 +/- 0.23 g/L of ethanol with 66.21 +/- 3.26% and 60.46 +/- 3.02% of fermentation efficiency, 0.33 +/- 0.02 and 0.31 +/- 0.02 g/g of ethanol yield and 0.24 +/- 0.01 and 0.13 +/- 0.01 g/L/h of productivity, respectively. The co-culture of *S. cerevisiae* (VS3) and *P. stipitis* (NCIM 3498) helped to improve all parameters i.e. 10.94 +/- 0.53 g/L of ethanol with fermentation efficiency, ethanol yield and productivity of 83.11 +/- 0.42%, 0.420 +/- 0.01 g/g and 0.30 +/- 0.01 g/L/h, respectively.

## 1 Introduction

Lignocellulosic substrates are in great abundance, easily available, relatively low-cost and have potentiality to produce clean fuel when compared to sugar or starch containing feedstocks (Naseeruddin et al., 2013; Romani, Garrote, Ballesteros, & Ballesteros, 2013). Hence, exploitation of these sources may provide sustainable energy supply at local, regional and national level and thereby improving the economy of the nation (Balat & Balat, 2009) without hampering food security. Moreover, bioethanol has got the adaptability to existing engines and if used in transportation can replace 30% of gasoline use (Krishnan et al., 2010; Oleskowicz, Thomsen, Schmidt, 2011).

*Prosopis juliflora* is a perennial deciduous thorny shrub, commonly growing in semi-arid region of Indian subcontinent, Saudi Arabia and the United States of America (Gupta, Sharma & Kuhad, 2009). It is an invasive species and considered as threat to the ecosystem affecting adversely the biodiversity of the region. However, the tree is valued for the shade, forage and can tolerate drought, salinity as well as grazing (Pasha, Thabit, Kuhad, & Linga, 2008). To eradicate this tree species mechanical, chemical and biological control programmes are commonly employed but there are other options available to utilize positively for human welfare (Rilov & Crooks, 2009). The viable options to manage the menace of these invasive plants for human welfare include its use as fuel wood and application in biofuel industry (Rilov & Crooks, 2009) as its carbohydrate content is 69.25% (on dry weight basis). As the tree does not form a part of main food or feed cycle, it qualifies to be a suitable substrate for long-term sustainable production of bioethanol (Pasha, Thabit, Kuhad, & Linga, 2008).

It is recommended that prior to distillation the fermented broth should contain higher concentration of ethanol which makes it imperative that initial sugars in the hydrolysate should be high. Concentration of initial sugars in the water soluble fraction of hydrolysate can be achieved by evaporation of water or addition of less water to the hydrolysis process or concentrating the hydrolysate at 70°C under vacuum (Srilekha Yadav, Naseeruddin, Sai Prashanthi, Sateesh & Venkateswar Rao, 2011). Carvalho, Mussatto, Cândido, & Almeida e Silva, (2006) concentrated the acid hydrolysate of *Eucalyptus* shavings and reported proportional increase in the sugar content as a function of the concentration factor employed.

The hydrolysate obtained after acid saccharification of lignocellulosic biomass contains toxic compounds generated mainly due to the degradation of sugars from hemicellulose and cellulose. These inhibitors interfere with the physiology of yeast cells resulting in decreased cell viability, ethanol yield and productivity (Chandel, Kapoor, Singh & Kuhad, 2007; Hahn-Hagerdal, Karhumaa, Fonseca, Spencer-Martins, & Gorwa-Grauslund, 2007). To minimize the effect of these inhibitory compounds several detoxification methods have been studied, but the effectiveness of each method depends on the type of hydrolysate and microorganism used (Anish & Rao, 2009). Moreover, each detoxification method has specificity for certain compounds; therefore combining one or more methods can yield better results for detoxification of the acid hydrolysate (Carvalho, Mussatto, Cândido, & Almeida e Silva, 2006). Overliming followed by neutralization and charcoal treatment method is most widely used and is considered as a promising and economic method of detoxification (Pasha, Thabit, Kuhad, & Linga, 2008; Jun-jun, Qiang, Yong & Shi-Yuan, 2009; Chandel, da Silva & Singh, 2011a).

Choice of organism is important in ethanol production from lignocellulosic materials as the hydrolysate contains mixture of both hexoses and pentoses like dextrose, galactose, mannose, xylose and arabinose. For economical production of lignocellulose-based ethanol, simultaneous utilization of these biomass-derived sugars is required (Lin and Tanaka, 2006). Although many organisms are available, most of the microorganisms lack efficiency to ferment both hexose and pentose sugars present in the hydrolysate. Though, *Saccharomyces cerevisiae* and *Zymomonas mobilis* are most commonly used microorganisms for ethanol production, they lack the ability to utilize pentose sugars (Hector et al., 2011). *S. cerevisiae* is mostly used in industrial processes for ethanol production because of several advantages such as high ethanol productivity, high tolerant of high ethanol concentrations and inhibitors associated with it (Hickert, da Cunha-Pereira, de Souza-Cruz, Rosa, & Ayub, 2013). Other yeasts like *P. stipitis, Candida shehatae* and *Pachysolen tannophilus* have also attracted interest as choice of organisms as they can convert pentose sugars into ethanol but are less tolerant of ethanol and inhibitors and also require a small and well-controlled supply of oxygen for maximum ethanol production (Hickert, da Cunha-Pereira, de Souza-Cruz, Rosa, & Ayub, 2013). Despite many attempts, the microbial strains improved by genetic engineering or protoplast fusion showed preference for one sugar over another as a result of catabolite repression leading to longer time for the complete utilization and fermentation of all the sugars. Therefore, co-culture fermentation for the fermentation of mixture of sugars is a viable option (Srilekha Yadav, Naseeruddin, Sai Prashanthi, Sateesh, & Venkateswar Rao, 2011) which circumvents the problems associated with monoculture of the wild strains or engineered yeast strains (Hickert, da Cunha-Pereira, de Souza-Cruz, Rosa, & Ayub, 2013). Therefore, in the present study an attempt has been made to utilize concentrated hydrolysate of *P. juliflora* pre-treated with dilute acid where both the sugars can simultaneously get fermented to ethanol by co-culturing with *S. cerevisiae* (VS3) and *P. stipitis* (NCIM 3498).

## 2 Materials and Methods

### 2.1 Microorganisms

#### 2.1.1 Saccharomyces cerevisiae

VS3: is an isolate of our laboratory that originated from the soil samples collected within the hot regions Thermal Power Plant, Kothagudem India (Kiran Sree, Sridhar, Suresh, Banat, & Venkateswar Rao, 2000). It was maintained on yeast extract, peptone, dextrose and agar (YEPDA) containing 10, 20, 20 and 25 g/L, respectively. The pH of the medium was adjusted to 5.5±0.2.

#### 2.1.2 Pichia stipitis

(NCIM 3498) was obtained from National Collection of Industrial Microorganism (NCIM), National Chemical Laboratory, Pune, India and was maintained on media containing Malt extract (5 g/L), Glucose (5 g/L), Xylose (30 g/L), Yeast extract (5 g/L), Peptone (20 g/L) and agar-agar (25g/L). The pH of the medium was adjusted to 5.5 ± 0.2.

### 2.2 Concentration of hydrolysate to increase sugar concentration

Acid hydrolysate (1845 ml) obtained at 100 g level of substrate as reported earlier Shaik Naseeruddin, Suseelendra Desai and L Venkateswar Rao, 2017 was concentrated to increase the sugar concentration for ethanol fermentation by vacuum distillation (assembled in the lab). During the vacuum distillation, the boiling temperature of the liquid was maintained at 70°C as per Dehkhoda & Brandberg, (2008). Sugars, phenolics and furans were checked before and after the concentration process.

### 2.3 Detoxification of concentrated acid hydrolysate

The concentrated hydrolysate was detoxified by overliming with CaO, followed by pH adjustment to 6.00±0.5 using 6 N H_2_SO_4_ and treated with charcoal. Sugars, phenolics and furans were estimated before and at each step of detoxification process. The acid hydrolysate was first detoxified by CaO addition at room temperature till the pH reached 10.5±0.5 under fast stirring. The mixture was incubated for 1 h with intermittent stirring to precipitate the inhibitors. The slurry was then filtered under vacuum to get clear filtrate by removing precipitates as described by Chandel, Kapoor, Singh & Kuhad, 2007. The pH of the clear filtrate obtained after treating the hydrolysate with CaO was adjusted to 6.00±0.5 by using 6N H_2_SO4 and again filtered under vaccum to remove traces of salt precipitates as described by Chandel, Kapoor, Singh & Kuhad, 2007. After overliming and neutralization the hydrolysate was treated with 3.5% (w/v) of activated charcoal and stirred for 1 h as described by Martinez, Rodriguez, York, Preston, Ingram, (2000). The mixture was centrifuged at 3000 rpm for 20 min followed by filtration under vacuum to remove traces of precipitates.

### 2.4 Fermentation of pentoses and hexoses by mono- and co-culture

Concentrated hydrolysate was used as substrate for fermentation studies by performing mono- and co-culture studies. The organisms utilized for the studies were hexose utilizing strain *viz Saccharomyces cerevisiae* (VS3) and pentose utilizing strain *Pichia stiptis* (NCIM 3498) (Srilekha Yadav, Naseeruddin, Sai Prashanthi, Sateesh, & Venkateswar Rao, 2011).

#### 2.4.1 Inoculum preparation

Thermotolerant yeast *Saccharomyces cerevisiae* (VS3) inoculum was prepared by growing the microorganism on YEPD medium consisting of (g/L): yeast extract, 10; peptone, 20; dextrose, 20; pH: 5.5 ±0.2 for 48 h at 37±0.5°C and 100 rpm in Erlenmeyer flasks on a rotary shaker (Pasha, Kuhad, & Linga, 2007). This 48 h inoculum with an O.D of 0.6 (10%, v/v) at 600 nm prepared in 50 ml YEPD broth was transferred to fermentation medium (Gupta, Sharma & Kuhad, 2009; Chandel, Kapoor, Singh & Kuhad, 2007)

Inoculum of *Pichia stipitis* (NCIM 3498) was prepared by growing it aerobically in Erlenmeyer flasks on a rotary shaker in medium consisting of (g/L): Xylose - 5; dextrose - 5; yeast extract - 5; malt extract -5; peptone - 5; pH 5.5±0.2 for 24 h at 30 °C and 100 rpm. This 24 h old inoculum with an O.D. of 0.6 (10 %, v/v) at 600 nm was transferred to fermentation medium (Gupta, Sharma & Kuhad, 2009; Pasha, Kuhad, & Linga, 2007).

#### 2.4.2 Fermentation medium

The concentrated and detoxified acid hydrolysate was used for fermentation. Reducing sugars present in the hydrolysate were used as source of carbon hence no other external carbon source was added to the hydrolysates. The hydrolysate was supplemented with 0.1% (w/v) each of yeast extract, peptone, NH_4_Cl, KH_2_PO_4_; and 0.05% (w/v) each of MgSO_4_.7H_2_O, MnSO_4_.5H_2_O, CaCl_2_.2H_2_O, FeCl_3_.2H_2_O and ZnSO_4_.7H_2_O before fermentation. The pH of medium was adjusted to 5.5±0.1 and autoclaved at 110°C for 20 min (Pasha, Kuhad, & Linga, 2007), cooled to 30±2°C and used for fermentation.

#### 2.4.3 Ethanol production by monoculture with hydrolysate at 100g level

Monoculture studies with *Saccharomyces cerevisiae* (VS3) and *Pichia stiptis* (NCIM 3498) were carried out separately in individual flasks using detoxified and concentrated acid hydrolysate obtained at 100 g level of substrate. At first, 600 ml of detoxified concentrated hydrolysate was supplemented with nutrients. A 30 ml of hydrolysate was taken from 600 ml of hydrolysate in three different flasks for fermentation. The pH of the hydrolysate was adjusted to 5.5±0.1 and autoclaved, inoculated with 10% (v/v) inoculum of each organism separately. The shake flask fermentation was carried out at 30±2°C, 150 rpm for 72 h and samples were collected at regular intervals to estimate residual sugar and ethanol concentration.

#### 2.4.4 Ethanol production by co-culture with hydrolysate at 100g level of substrate

About 150 ml of hydrolysate was taken from 600 ml of detoxified and concentrated acid hydrolysate obtained at 100 g level of substrate and supplemented with nutrients. The pH of medium was adjusted to 5.5±0.1, autoclaved (Pasha, Kuhad, & Linga, 2007) and inoculated with co culture of *Pichia stipitis* (NCIM 3455) + *Saccharomyces cerevisiae* (VS3) (added one after the other with an interval of 18 h). The flasks were incubated on a shaker; samples were collected at regular intervals, centrifuged at 6000 rpm for 10 min at 4°C and analyzed for residual sugars and ethanol.

### 2.5 Analytical Methods

The total reducing sugars released during delignification and after acid hydrolysis were estimated by DNS method (Miller, 1959). The fermentation inhibitors viz. furans and phenolics were analyzed spectroscopy. Phenolics were estimated by Folin - Ciocalteus method (Singleton and Rossi, 1965), and furans by Martinez, Rodriguez, York, Preston, Ingram, (2000). For estimation of phenolics, to the test tube containing 0.1 ml of sample mixed with 8.4 ml of distilled water, 0.5 ml of Folin-Ciocalteus reagent was added. Later after 5 min, 1 ml of 10% sodium carbonate was added. A blank without sample served as control. The reaction mixture was incubated at room temperature for 1 h and read at 750 nm against blank with gallic acid as standard. For estimation of furans, the hydrolysate obtained was neutralized and read at 284 nm and 320 nm against water as blank. The furan concentration at 320 nm was subtracted from the concentration at 284 nm and a correction factor of 0.056 was added to the final value. Fufuraldehyde was used as standard for total furans estimation. Cell density was measured turbidometrically at 600 nm. Fermentation broth was diluted 10 times using sterile water and turbidity was measured with a UV–VIS spectrophotometer 117 (Systronics India).

Ethanol produced was analyzed by gas chromatography (Shimadzu GC-2011-Japan) using ZB-Wax column (30 mm x 0.25 mm) with a flame ionization detector. The analysis was performed according to National Renewable Energy Laboratory procedure LAP #001 (David, 1994). The column temperature was maintained at 150°C (isothermal) and the carrier gas nitrogen was run at pressure of 16 kPa with a run time of 5.5 min as ethanol has retention time of about 2.3 min. The injector temperature was at 175°C and the detector temperature was at 250°C. The other parameters used were flow rate: 40 ml/min, spilt ratio: 1/50, velocity of H_2_ flow: 60 ml/min, sample quantity: 1 μl. The sample used for ethanol estimation was filtered by 0.22 μm cellulose acetate filter for GC analysis.

### 2.6 Statistical Analysis

All the experiments were conducted in triplicate and mean and standard deviation (SD) values were calculated using MS Excel software. Anova was carried out by using SPSS statistical package software version 19.0.

## 3 Results

### 3.1 Concentration of acid hydrolysate

Acid hydrolysate obtained at 100 g level of substrate was concentrated to increase the sugar concentration by vacuum distillation. After concentration, the sugar concentration reached to 33.59±0.52 g/L from initial level of 18.70±0.16 g/L, representing about two-fold increase. The main aim of vacuum distillation of acid hydrolysate was to increase the concentration of sugars but phenolics content also got nearly doubled and reached 3.94±0.18 g/L from initial concentration of 2.16±0.10 g/L. The furans concentration increased to 1.72±0.12 g/L from initial level of 1.10±0.03 g/L with overall decrease in total volume of hydrolysate from 1845 ml to 922 ml. Concentrated hydrolysate obtained (922 ml) was detoxified by overliming, neutralization with sulphuric acid followed by charcoal addition. By this treatment, there was a reduction of 4.73% in sugar concentration i.e. from 33.59±0.52 g/L to 32.00±0.48 g/L, however, with significant decrease in phenolics content from 3.94±0.18 g/L to 0.558±0.11 g/L (85.83% decrease) and furans content from 1.72±0.12 g/L to 0.21±0.08 g/L (97.79% reduction) with total final volume decreasing to 600 ml **(Table 1)** at p<0.05 F (2, 6) = 1143.038, 472.644 and 238.935 for sugars, phenolics and furans respectively.

### 3.2 Ethanol production by monoculture with acid hydrolysate at 100 g level of substrate

A 60 ml of hydrolysate containing a sugar concentration of 32 ±0.48 g/L was taken from 600 ml of concentrated and detoxified acid hydrolysate obtained at 100 g level of substrate and distributed equally (30 ml) into two 100 ml Erlenmeyer flasks. The hydrolysate in each flask was supplemented with nutrients and inoculated with 10% (v/v) *Saccharomyces cerevisiae* (VS3), and *Pichia stipitis* (NCIM 3498) separately. *Saccharomyces cerevisiae* (VS3) produced ethanol of 8.52±0.43 g/L by consuming 25.62±1.28 g/L of sugars from initial sugar concentration of 32 ±0.48 g/L with fermentation efficiency, ethanol yield and productivity of 66.21±3.26%, 0.33±0.02 g/g and 0.24±0.01 g/L/h, respectively, **(Fig 1)**. The pentose sugar fermenting yeast *Pichia stipitis* (NCIM 3498) produced 4.52 ±0.23 g/L of ethanol by consuming 14.66±0.87 g/L of sugars with fermentation efficiency, ethanol yield and productivity of 60.46±3.02%, 0.31±0.02 g/g and 0.13±0.01 g/L/h, respectively, **(Fig 1)**.

**Fig. 1:**
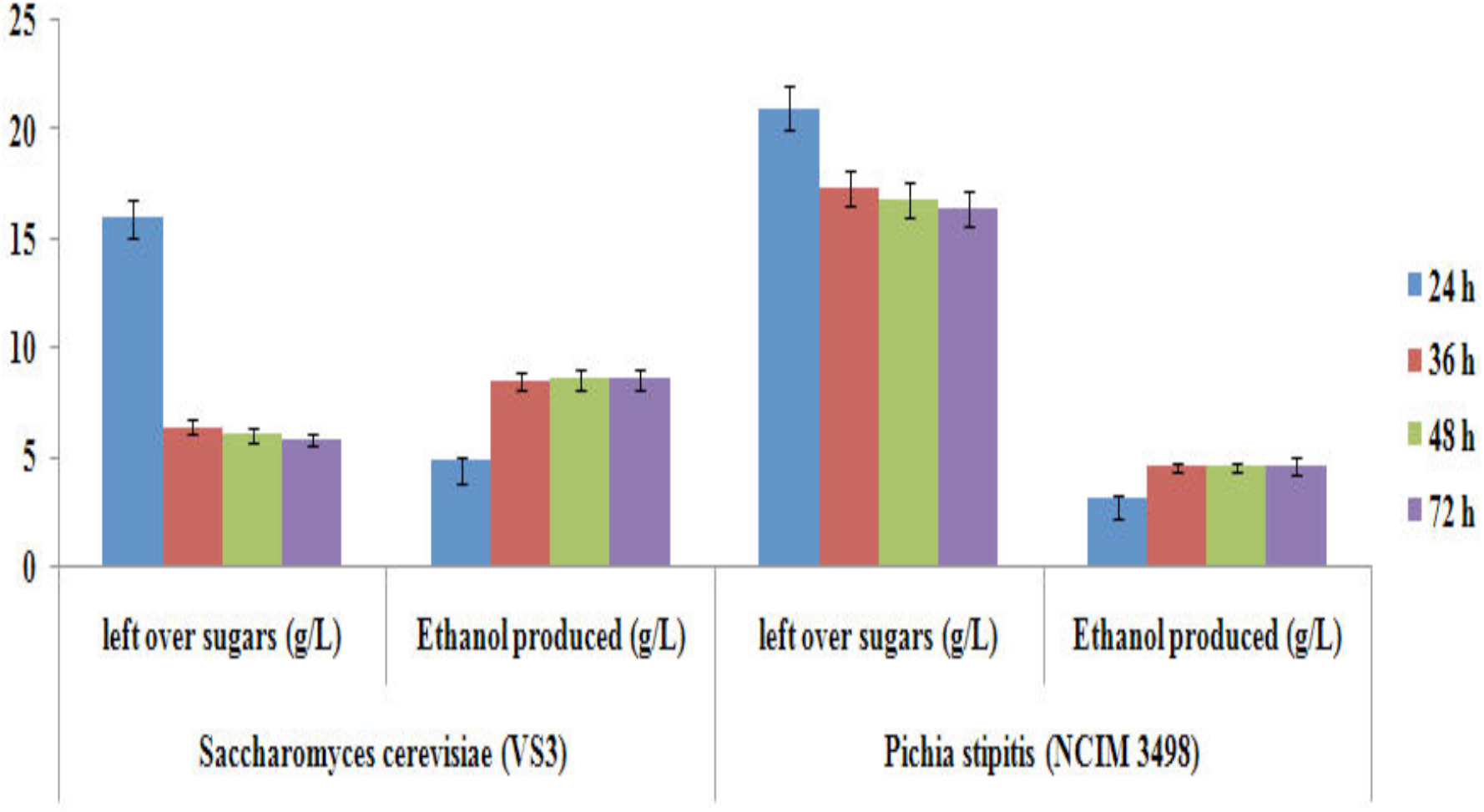
Fermentation of concentrated and detoxified acid hydrolysate (100g level) with two yeast strains.

### 3.3 Ethanol production by co culture with acid hydrolysate at 100g level of substrate

In co-culture with concentrated and detoxified acid hydrolysate (150 ml) obtained at 100 g level of substrate containing a sugar concentration of 32±0.48 g/L as substrate, addition of *Saccharomyces cerevisiae* (VS3) after 18 h of addition of *Pichia stipitis* (NCIM 3498) utilized 25.81±0.24 g/L of sugars and produced 10.94±0.53 g/L of ethanol with fermentation efficiency, ethanol yield and productivity of 83.11±0.42%, 0.420±0.01 g/g and 0.30±0.01 g/L/h, respectively **(Fig. 2)**.

**Fig. 2:**
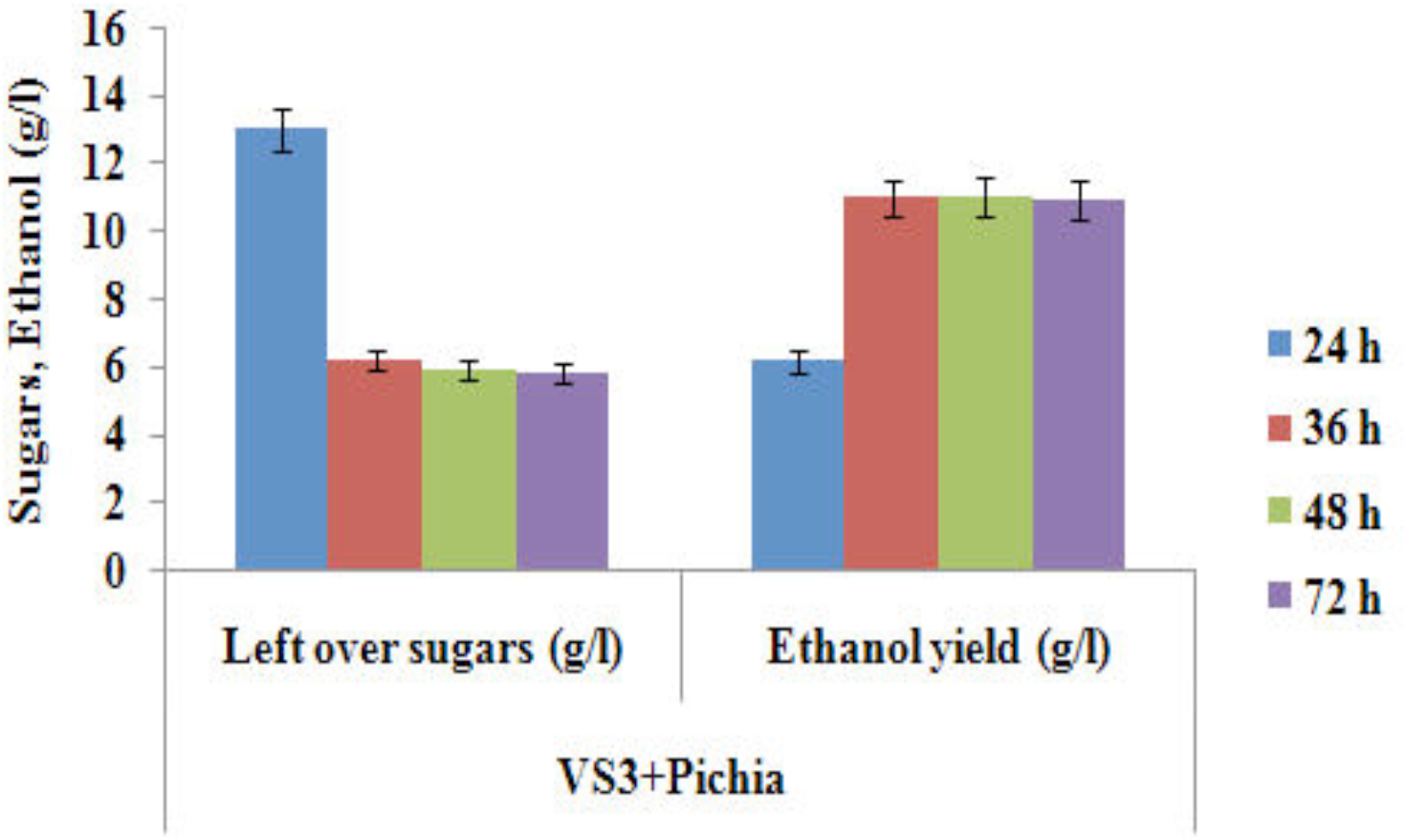
Fermentation of *Prosopis juliflora* concentrated and detoxified acid hydrolysate (100g level) with co culture of VS3+Pichia.

## 4 Discussion

Srilekha Yadav, Naseeruddin, Sai Prashanthi, Sateesh, & Venkateswar Rao, (2011) reported that batch fermentation of concentrated and detoxified acid hydrolysate of rice straw using coculture of *Saccharomyces cerevisiae* (OVB 11) and *Pichia stipitis* (NCIM 3498) resulted in maximum ethanol production after 36 h of incubation (12 g/L) with an efficiency of 95%. The volumetric ethanol productivity was 0.33 g/L/h with a yield of 0.4 g/g. In the present study also, 10.94±0.53 g/L of ethanol with fermentation efficiency, ethanol yield and productivity of 83.11±0.42%, 0.420±0.01 g/g and 0.30±0.01 g/L/h, respectively, was obtained. The increased ethanol yield and ethanol efficiency is due to the total conversion of both sugars i.e. dextrose and xylose. Chandel, Singh, Narasu & Rao, (2011b) performed the fermentation of hemicellulose acid hydrolysate with co-culture using *Pichia stipitis* (NCIM 3498) and thermotolerant yeast *Saccharomyces cerevisiae* (VS3) and reported 15.0 ±0.92 g/L of ethanol production with ethanol yield of 0.48±0.032 g/g and volumetric ethanol productivity of 0.208±0.0142 g/L/h. The findings of Sornvoraweat, Buddhiporn, Jirasak & Kongkiattikajorn, (2009) also support the present study who reported that ethanol yield was maximum from acid hydrolysate of *cassava* peels when cocultured with *Saccharomyces cerevisiae* and *Candida tropicalis* in comparison to monoculture of *Saccharomyces cerevisiae*.

## 5 Conclusions

The present study was carried out by utilizing concentrated *Prosopis juliflora* acid hydrolysate to ferment both the sugars simultaneously into ethanol by co-culturing of *Saccharomyces cerevisiae* (VS3) and *Pichia stipitis* (NCIM 3498) and able to produce 10.94±0.53 g/L of ethanol with ethanol yield of 0.420±0.01 g/g and volumetric ethanol productivity of 0.30±0.01 g/L/h. Results obtained in the current study suggest the feasibility of scaling up of production of ethanol from lignocellulosic acid hydrolysate of *Prosopis juliflora* for second-generation ethanol production co-culture of *Pichia stipitis* and *Saccharomyces cerevisiae* (VS3).

## 6 Acknowledgements

We thank the Department of Biotechnology (DBT), Ministry of Science and Technology, UGC-MANF and DRS-I SAP (Government of India) for the financial assistance.

